# A 3D printable device allowing fast and reproducible longitudinal preparation of mouse intestines

**DOI:** 10.1101/2021.12.01.470784

**Authors:** Beckey DeLucia, Sergey Samorezov, Megan T Zangara, Rachel L Markley, Lucas J Osborn, Karlee B Schultz, Christine McDonald, Jan Claesen

## Abstract

Accurate and reproducible analysis of mouse small and large intestinal lumen is key for research involving intestinal pathology in preclinical models. Currently, there is no easily accessible, standardized method that allows researchers of different skill levels to consistently dissect intestines in a time-efficient manner. Here, we describe the design and use of the 3D printed “Mouse Intestinal Slicing Tool” (MIST), which can be used to longitudinally prepare murine intestines for further analysis. We benchmarked the MIST against a commonly used procedure involving scissors to make a longitudinal cut along the intestines. Use of the MIST halved the time per mouse to prepare the intestines and outperformed alternative methods in smoothness of the cutting edge and general reproducibility. By sharing the plans for printing the MIST, we hope to contribute a uniformly applicable method for saving time and increasing consistency in studies of the mouse gastrointestinal tract.

## Introduction

In research, it is important to have uniform methods and practices to attain reliable, high-quality results within and across research institutes^1–4^. Histological evaluation of intestinal tissue is vital for assessing pathology in many different disease models. Consistent and uniform preservation of tissue samples allows for accurate assessment of biological replicates and easier comparison between multiple groups. For example, in fields utilizing animal preclinical models of colorectal cancer, the enumeration and measurement of murine intestinal adenomas provide critical data^5, 6^. The ability to open mouse intestines longitudinally to evaluate gross pathology of the intestinal lumen is therefore very important in gathering accurate adenoma data. While there is not an established standard method for dissection, researchers commonly use a pair of offset scissors to longitudinally cut open intestines, thereby revealing the lumen^7–11^. However, this difficult, time-consuming method leads to less than ideal visualization of adenomas^10^. Without cleanly dissected and well arrayed tissue, the accuracy of adenoma count and size could be compromised, leading to inaccurate data and varying results across studies performed by different research groups. Rudling *et al*. developed an alternative to the scissors method by constructing a “gut cutting” device from several pieces of metal^10^. This device consists of four blunt-end metal rods inserted through the lumens of the intestinal segments, which are then placed in a base unit. Next, a lid containing slanted cutting guides is placed on top and the intestines are manually cut with a scalpel along the guides. Utilization of this gut cutting device took significantly less time and resulted in higher quality preparation when compared to using scissors^10^. Later versions of this device were machined out of a solid block of duralumin^12^. Based on these later designs by Yoneda *et al*.^12^, we developed a similar, 3D printed version we call the “Intestinal Preparation Device” (IPD; Figure 1AB). In utilizing our IPD, we encountered a few drawbacks that underscored areas in need of improvement. Through multiple redesigns and trials, we engineered and optimized an easily 3D printed tool we named the “Mouse Intestinal Slicing Tool” (MIST; Figure 1CD). We propose that our 3D printable MIST provides an easily accessible and reproducible method to standardize the longitudinal dissection of mouse intestinal tissue across research groups and institutes.

**Fig. 1.**
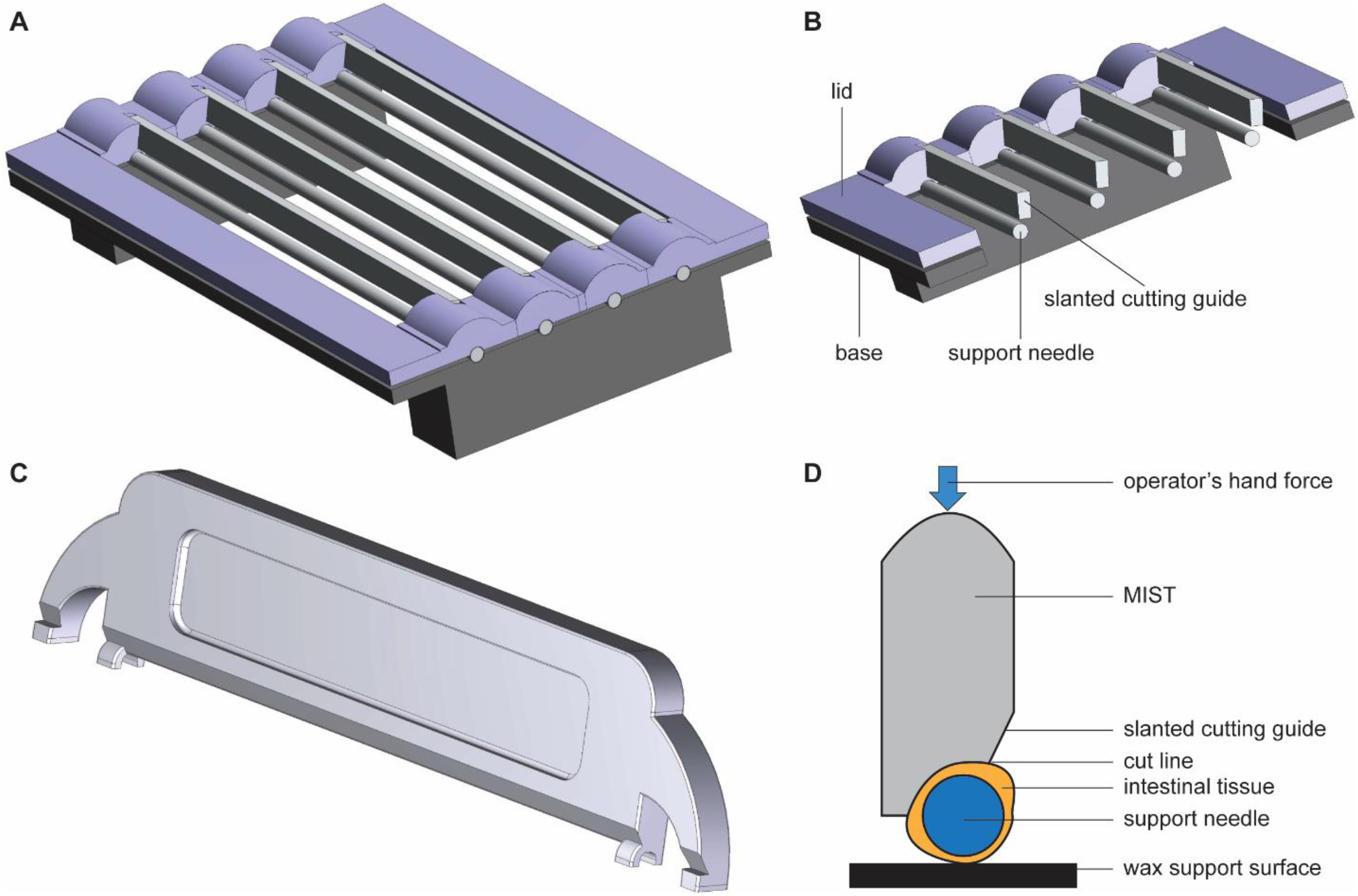
Schematic representations of the IPD and MIST. (A) Drawing of the IPD fully assembled with needles in place. (B) Cross section of the IPD showing the metal slanted cutting guides, which are used to tightly hold down the intestinal segment onto the support needle and are used to guide the scalpel during cutting. (C) Drawing of the MIST without needle, showing the forks that prevent the needle from rolling and bars at each end that prevent the needle from sliding longitudinally during cutting. (D) Schematic cross section of the MIST technique. The intestinal tissue (orange) is loaded onto the needle (dark blue) and kept in place between the device and the wax surface (black) through the operator’s downward hand force. The tissue is cut with a scalpel along the slanted cutting guide.

### Optimizing design of the IPD

To test the effectiveness and ease of various mouse intestine preparation methods, we utilized the *Apc*^*Min*/+^ genetic mouse model, which contains the Min (multiple intestinal neoplasia) mutant allele in its *Apc* (adenomatous polyposis coli) locus^5, 13^. This is a robust genetic model that predisposes mice to sporadic adenoma formation in both the small and large intestine.

To improve upon the widely used Scissors method (Figure 2AB), we initially used the IPD, our 3D printable version of the “gut cutting” device by Yoneda *et al*.^12^. Our IPD consisted of a base unit with indented wells for holding four metal support bars (knitting needles) that were previously inserted through the intestinal lumen and a lid with slots for detachable aluminum cutting guide bars (Figure 1AB). Utilizing 3D printing simplified the construction of the IPD in comparison to both versions of the gut cutting device, which were machined out of a series of metal plates or a duralumin block^12^. In addition, we utilized double pointed knitting needles for tissue stabilization, which offer an advantage over metal rods with rounded off ends. The tapered and rounded needle end makes insertion into the lumen easy, with a lower risk for creation of holes in the intestinal tissue (Figure 2CD). In using the IPD, we identified the need for different needle diameters to accommodate the narrowing lumen diameter along the intestinal tract to the colon (Figure 2EH). Too large a needle diameter distorted and/or created tears in the intestinal tissue. Conversely, if the needle diameter was too small, the tissues were not held securely in place during cutting. Another downfall of the IPD was encountered during transferring of the needles from the device onto the working surface (Figure 2H). Tissues could fall off the needle, making it difficult to smoothly array the intestinal lumen on our working surface. Moreover, the design of the IPD required that all four intestinal segments be threaded onto needles and cut open at the same time, meaning the last segment to be transferred onto the working surface was significantly drier and was more difficult to handle. Notably, despite the metal slanted guide bars, we found it difficult to cut in a straight line using the IPD due to the absence of a supporting surface directly underneath the needles. This led to the needles bowing down in the middle when pressure was applied. For this reason, we were not able to properly secure the tissues with the metal slanted cutting guide, resulting in jagged cut edges. Lastly, it was inconvenient and time consuming to assemble and dissemble the IPD.

**Fig. 2.**
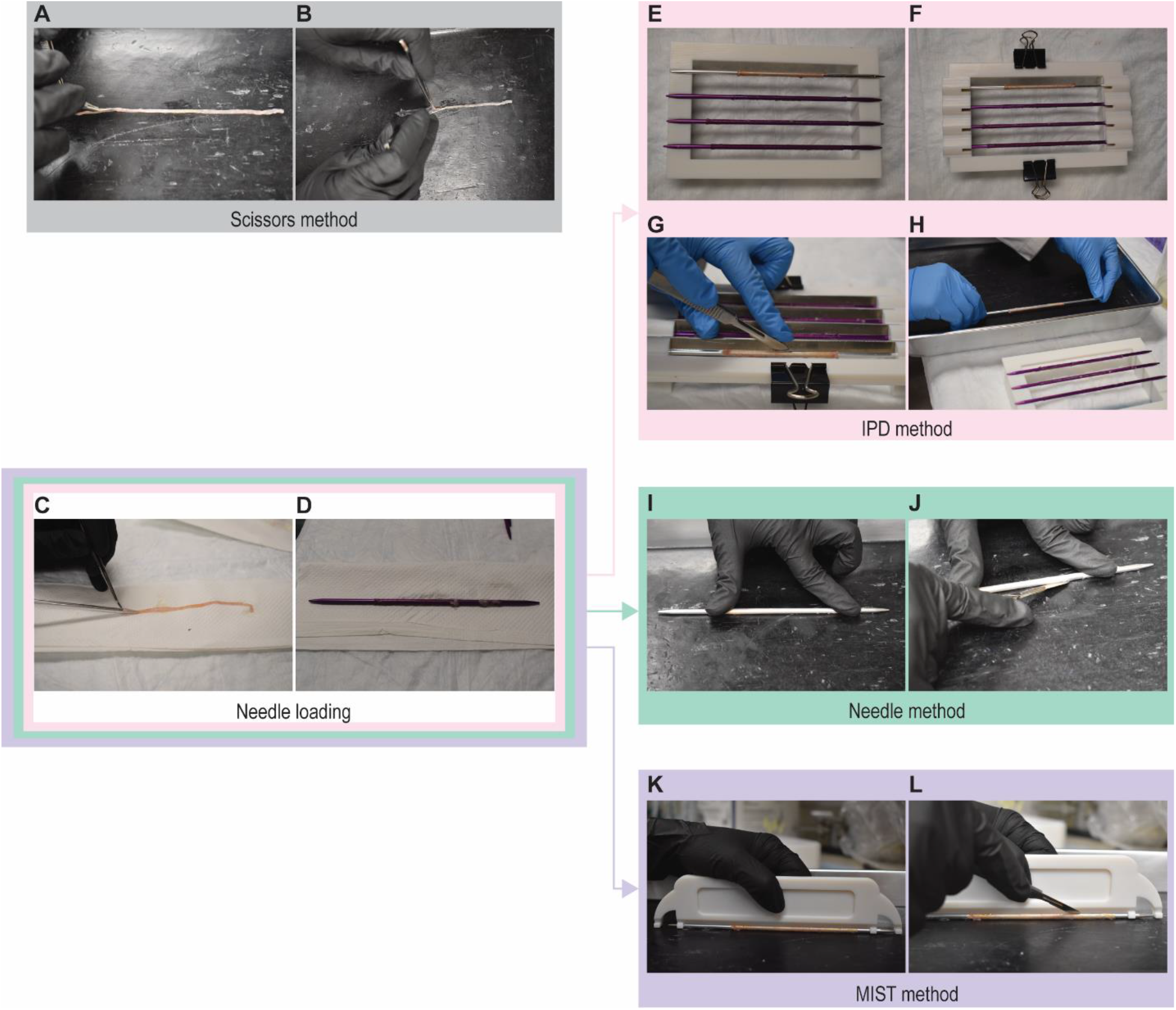
Overview of intestinal preparation methods. (A-B) **Scissors method** (A) The intestinal segment was cut open longitudinally one to two centimeters at a time using a pair of scissors. (B) The lumen was spread open using tweezers. (C-D) **Needle loading**, the initial step in the IPD, Needle, and MIST methods was inserting a needle through the lumens of the segments. (E-H) **IPD method** (E) loaded needles were placed into the base of the IPD and (F) secured in place by the lid and binder clips. (G) Metal slanted cutting guides were inserted into the lid and used to cut the tissue. (H) The device was disassembled, the loaded needles with cut intestines were transferred to the working surface and spread open. (I-J) **Needle method** (I) Tissue is secured at the proximal and distal ends of the segment by two fingers pressing the loaded needle against the working surface. (J) A scalpel is run down the length of the needle cutting the intestines. (K-L) **MIST method** (K) Tissue is secured in place uniformly by placing the MIST on top of the loaded needle and applying pressure with your hand. (L) The built in slanted cutting guide of the MIST is used to safely cut the intestine.

To prevent the needle holding the intestine from bending, we next tried a “Needle method” (Figure 2IJ). This method consists of pressing the intestine and needle against a wax dissection board which provides a support surface. Then a scalpel can be run down the length of the side of the needle. This removed the need for various device parts and no longer required transferring of tissues from a device to the working surface compared to the IPD. The largest drawback of this method is the pronounced lack of a safety guard between the operator’s fingers holding the needle and the scalpel blade. Additional shortcomings of this method were the lack of a cutting guide to allow for a straight cut, poor visualization of the intestine, and uneven pressure along the length of the intestine making the Needle method technically difficult.

Taking the aforementioned needs for improvement across the previous intestinal preparation devices into consideration, we engineered the MIST-a small, lightweight, cost-effective tool made with a 3D-printer. The MIST allows for easy use where one hand comfortably presses the MIST against the dissection tray, sandwiching the intestinal tissue in place with uniform pressure (Figure 1D, Figure 2KL). To accommodate the varying needle sizes required per section of the intestine, the MIST’s dimensions can be quickly adjusted by altering the printing plans. For example, we developed two variants of the MIST. One version has dimensions compatible with needle diameters of 3.50 to 3.75 mm and the other model that fits needle diameters of 2.75 to 3.25 mm. At both ends of the device, we included bars to prevent the needle from sliding out horizontally during the cut (Figure 1C). A built-in slanted cutting guide was also incorporated to permit for safe cutting with a scalpel by acting as a guard between the operator’s hand and the scalpel blade. The design permits the use of the device by either right- or left-handed individuals. Additionally, the design of the MIST allows for clear visualization of the cutting surface, leading to clean cut lines. Since the MIST does not contain excess crevices, it is easily cleanable with ethanol, surgical instrument cleaner, or disinfectants. The simple, small nature of the device also allows for ease of use in biosafety cabinets. Therefore, the MIST has the advantages of a simple design, improved tissue stabilization, and enhanced safety features over previous tissue dissection devices.

### The MIST preparation method consistently requires less time

We performed objective measurements of cut time and cutting edge accuracy to compare the performance of the MIST to the IPD, Needle, and Scissors dissection techniques. For mouse necropsies that involve analysis of both the small intestine (typically analyzed in three segments) and large bowel (analyzed as a single segment), the total amount of harvest time per mouse can quickly add up when using large experimental groups. Hence, we compared the amount of time required to longitudinally prepare the small and large intestines per mouse using the four different techniques (Figure 3). We found that the MIST, IPD, and Needle methods were all significantly quicker than the benchmark Scissors method, which took an average time per mouse of 12.2 minutes. The IPD method decreased the average time per mouse to 7.7 minutes. The Needle and MIST method further decreased the preparation time by roughly 50% with averages of 6.1 minutes and 6.2 minutes, respectively. In addition to the significant improvement in timing, the MIST method yielded the smallest range of preparation times, indicating good reproducibility between samples.

**Fig. 3.**
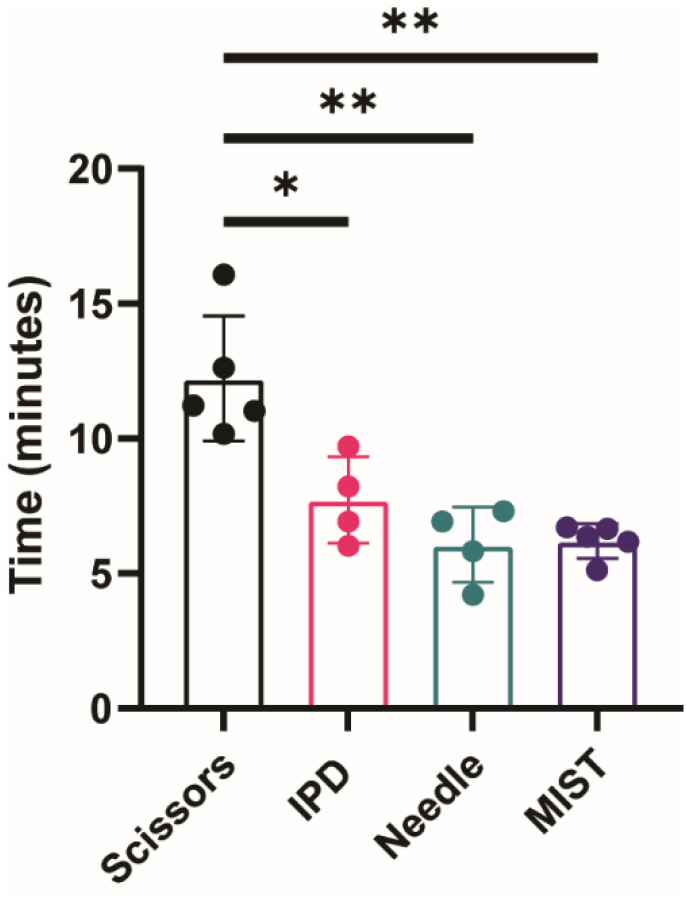
Comparison of preparation times across different methods. The time to longitudinally prepare the four intestinal segments per mouse was measured for the different preparation methods. While the preparation times when using the IPD were only slightly improved compared to the Scissors method (^*^*P* =0.0111), both the Needle and MIST methods had greatly improved prep times (^**^*P*=0.0020 and ^**^*P*=0.0032, respectively). Statistical analysis was performed using Brown-Forsythe and Welch ANOVA tests uncorrected for multiple comparisons. N= 4-5 per group.

### The MIST provides increased quality of intestine preparation

The resulting quality of the intestinal preparation using the various devices is visually evident (Figure 4AD). We noticed that the Needle (Figure 4C) and the MIST methods (Figure 4D) have smoother, straighter cut edges, while the Scissors (Figure 4A) and IPD methods (Figure 4B) yield many curves and lumps along the cut edge. To quantify this observation, we determined the ratio between the total segment length (measured along the middle of the tissue), and the length of the bottom cut edge (Figure 4E). A ratio of one represents a ‘perfect cut’, meaning the cut edge length is equal to the actual length of the segment, a greater ratio indicates a longer cut edge than the segment length. Both the Needle and MIST methods yielded ratios closer to one and were significantly lower than the benchmark Scissors method. Similar to the timing data, the MIST method had a tight range of experimental measurements.

**Fig. 4.**
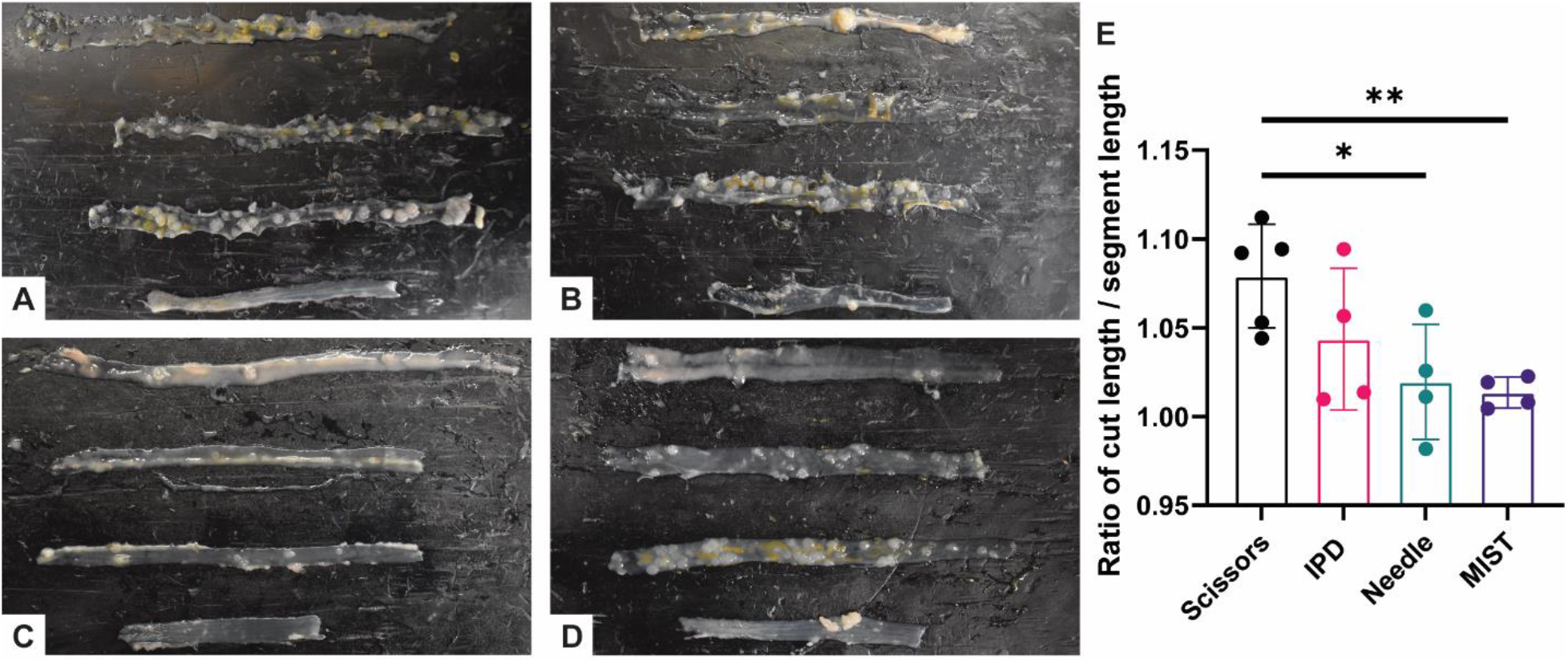
The MIST method reproducibly yields the neatest cutting edges. Representative photos of longitudinal intestine preparation using the (A) Scissors method, (B) IPD method, (C) Needle method, and (D) MIST method. The topmost segment is SI-1, followed by SI-2, SI-3, and Colon at the bottom. (E) The neatness of the cutting edge was compared to the actual segment length for each preparation method. A ratio of one represents a ‘perfect cut’, meaning the cut edge length is equal to the actual length of the segment. Compared to the Scissors method, the cutting edge quality was significantly improved for the MIST (^**^*P*=0.0054) and Needle (^*^*P*=0.0277), but not for the IPD (*P*=0.1915). Statistical analysis was performed using Brown-Forsythe and Welch ANOVA tests uncorrected for multiple comparisons. N= 4-5 per group.

### The MIST device allows for high quality Swiss-roll preparation for histology

The dimensions of the small and large intestine make it difficult to preserve in its native form, therefore the Swiss-roll technique was created as a method to preserve the integrity of large lengths of intestinal tissue for histological analysis^14^. This preparation allows for the visualization of the entire length of the mouse small or large intestine on one slide. The Swiss-roll technique is a straightforward method in which a longitudinally opened section of intestinal tissue is rolled in upon itself around a stick-like implement (toothpick or pin) prior to fixation. The resulting sample, once embedded, gives an uninterrupted, lateral view of the entire length of embedded tissue (Figure 5AC). Proper alignment of the tissue edges is important for creating a neatly rolled tissue sample, and aids in optimal orientation of tissue structure for histological analysis. When compared to colonic Swiss-roll samples cut using the Scissors method (Figure 5AB), MIST method-prepared Swiss-rolls were not only easier to roll, but also resulted in better crypt orientation (Figure 5CD). The even edge created with the MIST method decreased instances of rolled sample edges, allowing for more consistent sample orientation without the need to cut deeply into the paraffin block.

**Fig. 5.**
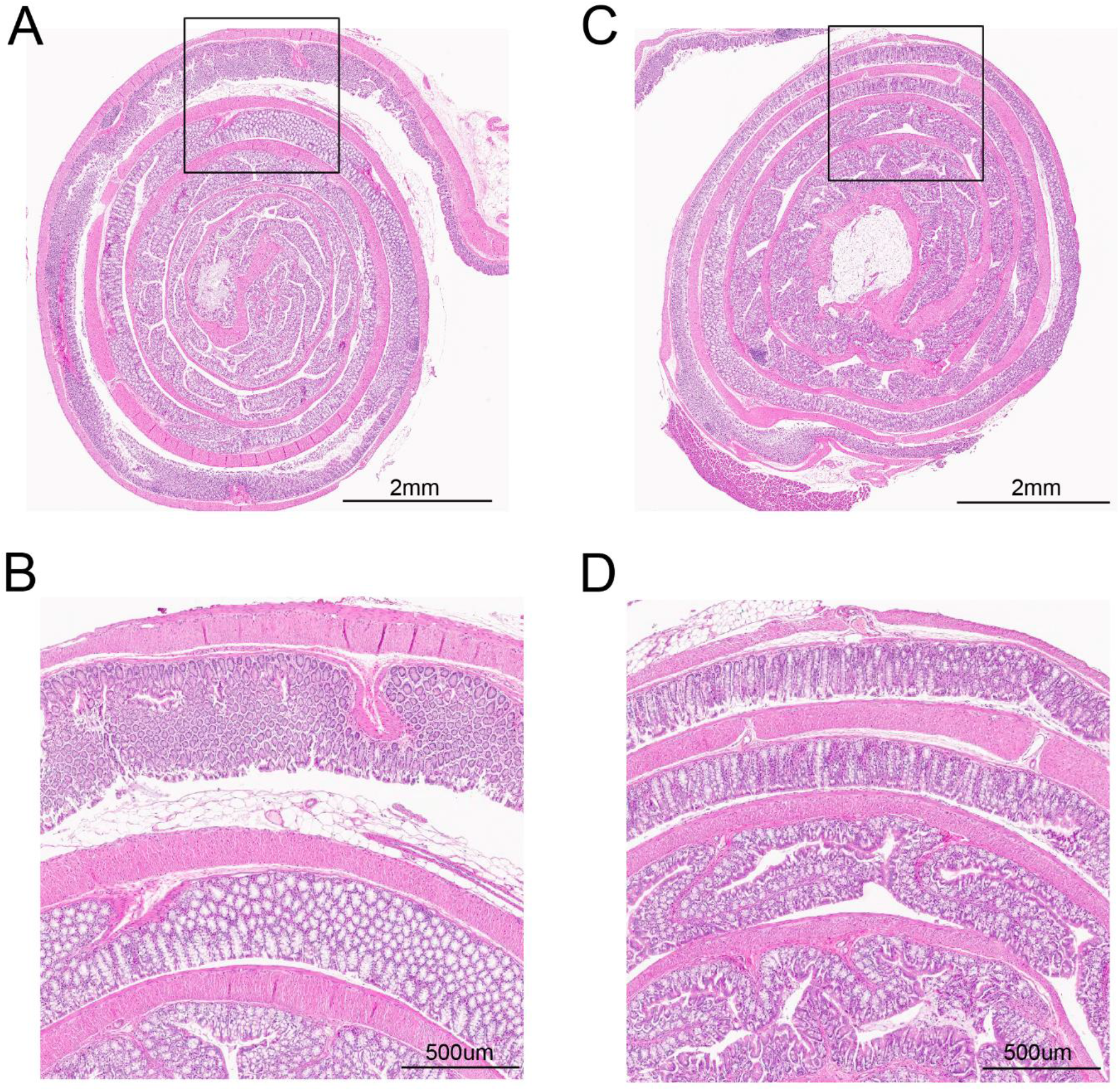
The MIST device allows for higher quality Swiss-roll histology. Representative H&E-stained colonic Swiss roll preparations using the (A-B) Scissors method and (C-D) MIST method. The insets (B and D) give a higher magnification view of the crypt orientation achieved by each method.

## Discussion

To achieve experimental data that can be reproduced with high integrity, is essential to validate discoveries and help advance our knowledge. It is crucial to have uniform practices to attain reliable, high-quality results within and across institutes. These techniques should be easy, replicable, convenient, and efficient to allow researchers of all skill levels to work with ease. Here, we have engineered and enhanced the design of the 3D-printed MIST, which we propose as a tool for providing simple, straightforward, and reproducible longitudinally cut mice intestines. To assess the efficacy of the MIST, we benchmarked it against the widely used Scissors method and compared it to two additional device-assisted methods, IPD and Needle. We objectively quantified the effectiveness of these methods by measuring the amount of time it took to prepare intestines and the straightness of the cut edges.

In measuring the amount of time to prepare the four intestinal segments per mouse, we showed that all experimental methods were significantly faster than the conventional Scissors method. Using the Scissors method, the intestines were cut and spread open a couple of centimeters at a time because if a segment was cut all at once it was both challenging and time consuming (due to the small, fragile tissue) to find the correct cut edges to neatly spread open the lumen. Using the Needle and MIST methods, we were able to prepare the intestines in half the amount of time, on average, compared to the Scissors method (Figure 3). This is a substantial advantage for experiments involving a large number of animals. For every ten mice that require intestinal preparations, using the MIST will save an average of 60 minutes. This time can be utilized toward exploring more research questions or performing additional experiments. In summary, the MIST provides a more time-efficient means of longitudinally preparing mice intestines.

The resulting quality of the intestinal preparations was visually assessed across the various methods. We noticed an appreciable difference between the intestine preparations using the Scissors and IPD methods versus the Needle and MIST methods (Figure 4AD). Based on visual observations, the Scissors and IPD resulted in very rough edges, obstructing researchers from having a clear view of the lumen for accurate adenoma enumeration or other macroscopic changes. For the Scissors method this was likely due to the lack of any cutting guide. Regarding the IPD method, the intestinal preparations had less smooth cut edges possibly due to the absence of a working surface directly underneath the loaded needles and the required transfer of the loaded needle from the device to the working surface. Although both the Needle and MIST methods resulted in clean, smooth cut edges, we observed that occasionally the Needle method resulted in a thin layer of intestine being cut off as seen on the SI-2 segment (Figure 4C). This was due to the lack of a cutting guide, poor visualization of the tissue, and having to repeatedly run the scalpel down the length of the needle simply due to the difficulty of trying to run a scalpel straight against a round needle. To corroborate our visual observations with objective measures, we quantified the ratios between the bottom cut edge to the actual segment length. From this, we showed that the Needle and MIST methods resulted in significantly straighter cut edges compared to the Scissors method, thereby upholding our visual observations (Figure 4E). In both of our objective measures, the Needle and MIST techniques had similar values. Despite these similar outcomes, using the MIST is more ideal for its consistency and safety. Considering both visual observations and objectively measured values, we conclude that the MIST method resulted in the highest quality preparation.

An important feature of any research technique is reproducibility. The MIST yielded the most constant preparation times across mice (Figure 3). Equally important, the MIST method produced the most consistent neat cut edges, as observed from its narrow range of values (Figure 4E). All things considered, it can be appreciated that the MIST is a highly dependable method for repeatedly preparing mice intestines of the same high caliber.

To test the applicability of the high-quality intestine preparation provided by the MIST, we formed Swiss rolls from colons cut open using Scissors or the MIST and submitted the preparations for hematoxylin and eosin staining (Figure 5). The even edge obtained by using the MIST resulted in better and more consistent tissue orientation when the blocks were sectioned at the same depth. This will normalize histological sample integrity for easier comparisons, as well as reduce waste of experimental samples. The high-quality preparations resulting from the use of the MIST can also be a powerful tool in making accurate gross histological observations. In the field of cancer research, the accuracy of the enumeration and measurement of adenomas can be improved using MIST. Additionally, researchers interested in inflammatory changes in the intestine can benefit from use of the MIST to visualize gross morphological changes indicative of inflammatory bowel disease. Ultimately, the MIST can be widely utilized across many diverse research niches.

In designing the MIST device, we sought to not only improve upon the reproducibility and efficacy of other previously established methods, but to also make it cost effective and easy to procure. The use of 3D printing in the fabrication of the device makes it easily reproducible and cost efficient. The device’s “.STL” and design files are sharable and modifiable, allowing for easy alteration to fit a range of research parameters. Additionally, the minimal parts required makes the MIST quick, safe to use, and easy to disinfect; for applications in a germ free setting, the MIST could be 3D printed using a resin that can withstand autoclaving.

In summary, given our visual and objective measures, we support the MIST as a strong candidate for a standard technique to achieve high quality longitudinal intestinal preparations for application in various research areas. In sharing the printing plans for the MIST we aspire to greatly enhance the resulting data for preclinical models of gastrointestinal studies.

## Methods

### Design of the IPD and MIST devices

The IPD device was thought of as an efficient way of handling multiple samples at one time by having four support needles loaded into the device (Figure 1). The shortcoming of this method was that the sample tissue was fixed/held only on the top (between the top surface of the supporting needle and the bottom surface of the cutting guide) but not at the bottom. In addition, the need to provide secure clamping of all four needles required increased fixture rigidity, leading to increase in the fixture weight. The MIST design was conceptually based on the idea that a needle with an intestine sample can be secured between the wax support surface at the bottom and the MIST on the top (Figure 1). The operator’s hand provides the well-controlled clamping force, while the sample to be cut is held in place both from the top and from the bottom. Semi-circular needle holders at the bottom of the MIST secure the needle radially while additional limiters on the MIST outside provide for axial stability. The MIST device as described in this paper has been 3D printed using Stratasys Vero material (https://www.stratasys.com/materials/search/vero). Other materials could be used given they satisfy the researcher’s requirements for cleanliness or sterility.

### Animals

All animal procedures were approved by the Cleveland Clinic Institutional Animal Care and Use Committee. Male C57BL/6 *Apc*^*Min*/+^ mice (C57BL/6J-Apc^*Min*^/J, stock#002020, Jackson Labs) and *Nod2-/-* mice (B6.129S1-Nod2^tm1Flv^/J, stock#005763, Jackson Labs) were housed under specific pathogen-free conditions and fed a standard breeder diet (Envigo Teklad Global Irradiated Rodent Diet 2018) in the Biological Resources Unit within the Cleveland Clinic Lerner Research Institute, Cleveland, OH. Mice between 5-6 months of age were euthanized and intestinal tissue excised for device testing.

### Preparation of intestinal segments

The small intestine was cut into three equal segments (proximal segment, mid segment, and distal segment) and referred to as SI-1, SI-2, and SI-3 respectively. SI-1, SI-2, SI-3, and the colon (C) were placed in a 150mm diameter petri dish containing 0.9% saline. Luminal contents of the intestinal segments were removed by flushing with 0.9% saline. Cleaned intestinal segments were lined on a black wax dissection tray for assessment of longitudinal opening by the four different methods described below. Timing of each method stopped once intestinal segments were spread open longitudinally on our working surface. Intestinal preparations were photographed for further analysis.

### Scissors Method (Figure 2AB)

1. Starting with SI-1, the intestinal segment was placed on a sheet of paper towel to remove excess saline. This allowed the tissue to stay in place on the working surface.
2. The tissue was placed onto the working surface vertically such that the proximal end was closest to the operator. Starting at the proximal end, one to two centimeters of the intestinal segment was cut using a pair of sharp-ball tip spring scissors (Fine Science Tools, Item No. 15033-09) (Figure 2A).
3. Using tweezers, the inner lumen was revealed by carefully pulling the cut edges apart (Figure 2B). The intestines were cut and spread open a couple of centimeters at a time because it was challenging and time consuming to find the edges and neatly spread open if the segment was cut all at once.
4. Steps 1 through 4 were repeated until all segments were laid open with the lumen exposed (Figure 4A).

### Needle loading (Figure 2CD)

The Needle, IPD, and MIST methods all required Needle loading as the initial step. Needle loading consisted of using a pair of tweezers to lift open the lumen and then inserting a needle through the lumen (Figure 2C) until the needle filled the length of the intestinal segment (Figure 2D). The needles used were aluminum knitting needles, double point (7 inches long, diameter Size 2-Size 5, Yarnology, MN) and were placed in 0.9% saline prior to loading allowing them to easily through the length of the segment. Once on the needle, tissue remnants on the outside of the intestine was carefully removed with scissors. The lumen size of the intestinal segments decreased as we went distally from the stomach to the anus. Hence, a variety of needle diameters were used depending on the diameter of the intestinal segment. For SI-1 we used needles with diameters of 3.75mm (Size 5) or 3.50mm (Size 4). For SI-2 and SI-3, needle diameters of 3.50mm (Size 4) or 3.25mm (Size 3) were used. With the lumen of the colon being the smallest, the needle diameters used were 3.25mm (Size 3) or 2.75mm (Size 2). Since the appropriate needle diameter to use is dependent on the size of the lumen, the most appropriate needle diameter to use may vary based on multiple factors. For example, mice model, sex, age, size, and treatment (inflammatory conditions) may increase or decrease lumen diameter.

### IPD Method (Figure 2EH)

1. All four intestinal segments were loaded onto needles.
2. The loaded needles were placed in the designated half-circle wells in the base of the IPD (Figure 2E). To maintain consistent orientation, the proximal end of each intestinal segment was loaded on the left side.
3. The lid was placed over the base containing the loaded needles. The base and lid are clamped together securing the needles in place horizontally and vertically (Figure 2F).
4. The 4 metal slanted cutting guides were inserted into the designated slots on the lid.
5. One hand was used to press the metal slanted cutting guide against the needle to secure the tissue in place. With the other hand, a scalpel was used to longitudinally cut the length of the segment (Figure 2G). This was repeated until all segments had been cut.
6. Disassembly of the device was achieved by carefully removing the metal cutting guides, unclamping, and removing the lid.
7. Starting with SI-1, the cut intestine around the needle was transferred to our working surface (Figure 2H). Then, using a gloved finger, the intestine was gently removed from the needle.
8. Step 7 was repeated until all segments were laid open with the lumen exposed (Figure 4B).

### Needle Method (Figure 2IJ)

1. All four intestinal segments were loaded onto needles.
2. The loaded needle containing SI-1 was placed on the working surface vertically with the proximal end closest to the operator. With one hand, the tissue was secured against the working surface and needle. The proximal end of the tissue was held by applying pressure with the tip of the pointer finger. At the distal end of the intestine, pressure was applied with the thumb. Significant pressure was applied to ensure the intestines did not slide on the needle while cutting (Figure 2I).
3. With the free hand, using a scalpel, a longitudinal cut was carefully made along the length of the needle. Extreme caution was exerted to avoid cutting fingers (Figure 2J).
4. Using a gloved finger, the intestinal segment was gently removed from the needle and spread open on the working surface.
5. Steps 2 through 4 were repeated until all segments were laid open with the lumen exposed (Figure 4C).

### MIST method (Figure 2KL)

1. All four intestinal segments were loaded onto needles.
2. The loaded needle containing SI-1 was placed onto the working surface vertically with the proximal end closest to the operator.
3. With one hand, the MIST was placed on top of the loaded needle. Pressure was evenly applied onto the tissue in all areas from the force of the hand pressing the MIST down (Figure 2K).
4. Using the MIST’s built-in cutting guide, a scalpel was used to longitudinally cut open the intestine (Figure 2L).
5. The MIST was removed and with a gloved finger, the intestine was gently removed from the needle.
6. Steps 2 through 5 were repeated until all segments were laid open with the lumen exposed (Figure 4D).

### Intestinal segment and cutting edge measurements

Measuring the neatness of the cutting edge in comparison to the middle or actual length of the intestines was achieved through image analysis in ImageJ. First, the prepped intestines were photographed with a reference ruler in frame. With the image opened in ImageJ a scale of one centimeter was set by tracing the distance of one centimeter on the reference ruler in the photograph. Using the segmented line tool, the bottom cut edge of SI-1 was traced and measured. Next, using the same segmented line tool, the middle length of SI-1 was measured.

### Swiss roll preparation and histology

Excised and flushed colons from Nod2^-/-^ mice were opened longitudinally using either the Scissor or MIST method and laid flat on the dissecting surface. The handle end of a sterile cotton swab was placed across the proximal end of the tissue and used as an anchor to roll the tissue around itself. Once fully rolled, the tissue roll was gently pushed off the end of the handle of the cotton swab using forceps into a single-chamber cassette. Rolls were fixed in Histochoice® Tissue Fixative (VWR) for 24 hours. After fixation, samples were paraffin embedded in which 5µm sections were cut, mounted on glass slides, and stained with hematoxylin and eosin. Slides were scanned into electronic files using an Aperio AT2 slide scanner at 20x magnification for histological evaluation.

## Supporting information

3D rendering of MIST for large needle

.STL file of MIST for large needle

3D rendering of MIST for small needle

.STL file of MIST for small needle

## Funding

The study was supported by a Research Grant from the Prevent Cancer Foundation (PCF2019-JC) and seed funding from the Cleveland Clinic Foundation (JC). JC is additionally supported by a National Institutes of Health grant (R01 AI153173), an American Cancer Society Institutional Research Grant (IRG-16-186-21) and a Jump Start Award (CA043703) from the Case Comprehensive Cancer Center.

## Author contributions

Conceptualization and design: BD, SS, JC. Investigation: BD, SS, MTZ, RLM, LJO, KBS, CM, JC. Figures: BD, SS, MTZ, CM, JC. Writing of the original draft: BD. Writing, review and interpretation: all authors.

## Competing interests

JC is a Scientific Advisor for Seed Health, Inc.

## Materials and correspondence

Design and .STL files for the MIST are freely available as supplementary information included with the article preprint. Please address all correspondence and material requests to JC at

